# Long-term memory in frog-eating bats

**DOI:** 10.1101/2022.03.16.484498

**Authors:** M. May Dixon, Patricia L. Jones, Michael J. Ryan, Gerald G. Carter, Rachel A. Page

## Abstract

Long-term memory has clear advantages but also has neurological and behavioral costs^1–3^. Given these opposing selection pressures, understanding how long memories last can shed light on how memory enhances or constrains animals’ abilities to exploit their niches. Although testing memory retention in wild animals is difficult, it is important because captive conditions do not reflect the complex cognitive demands of wild environments, and long-term captivity changes the brain^4^ (past studies on nonhuman long-term memory are reviewed in Table S1). Here, we trained wild-caught frog-eating bats (*Trachops cirrhosus*) to find prey by flying to a novel acoustic cue, released them back into the wild, and then re-captured some of them 1-4 years later. When re-tested, all eight ‘experienced’ bats that previously learned the novel prey sounds flew to those sounds within seconds, whereas 17 naïve bats tested with the same sounds showed weak responses. Experienced bats also showed behavior indicating generalization of memories between novel sounds and rewards over time. The frog-eating bat’s remarkably long memory for novel acoustic cues indicates that an ability to remember rarely encountered prey may be advantageous for this predator, and suggests hitherto unknown cognitive abilities in bats.

## Introduction and methods

The phyllostomid bats are the most extensive adaptive radiation of any mammalian family within the most ecologically diverse mammalian order. The predatory phyllostomid *T. cirrhosus* is an emerging model in cognitive ecology^5,6,8^, which hunts by eavesdropping on the mating calls of many frog and katydid species, and can discriminate the calls of palatable versus poisonous species^5^. We captured 49 wild *T. cirrhosus*, individually marked them, and trained them to fly to a novel sound (one of two ringtones: “trained-A” or “trained-B”)^6^. After training, bats spontaneously generalized the association and flew to other ringtones. We then trained the bats to discriminate between their trained ringtone and three other unrewarded ringtones^6^. Before release, these ‘experienced’ bats had retrieved rewards in response to flying to their trained ringtone at least 40 times over 11 to 27 days.

We recaptured 8 of the 49 experienced bats 356–1531 days after their initial release, and retested them on their trained ringtone under the same conditions as their original training^6^. To investigate how much bats would generalize the response to similar stimuli, we also played an ‘extinguished ringtone’ and a ‘control sound’. The extinguished ringtone was one of the acoustically similar but unrewarded ringtones used in their discrimination training (ringtone “E” in ^6^). The control sound was an acoustically different 15-kHz pure tone, selected to assess whether bats would generalize the experimental association to any sound from the speaker. As a control group, we presented the same sounds to 17 wild-caught naïve bats with no prior experience with the experimental sounds. We scored the maximum responses of the naïve and experienced bats using an ordinal scale of increasingly strong responses: 0 = *no response*, 1 = *ears twitched in time with stimulus (see supplemental video)*, 2 = *approached stimulus within 1 m*, and 3 = *attacked speaker and retrieved reward*.

## Results

When played the trained sounds, the experienced bats clearly responded more than the naïve bats (Fig. 1, permutation test: P<0.0002, see supplement). For example, six of eight experienced bats attacked the trained ringtone and all eight approached it, whereas none of 17 naïve bats attacked and only one approached (Fig. S1). The experienced bats’ responses to the trained and extinguished sounds were stronger than to the control sounds (P=0.001, Fig. 1), whereas the naïve bats typically only twitched their ears to all sounds (Fig. S1). We saw no clear evidence that the experienced bats’ responses decreased across the retention times of 356 to 1531 days (Fig. S1).

**Fig. 1.**
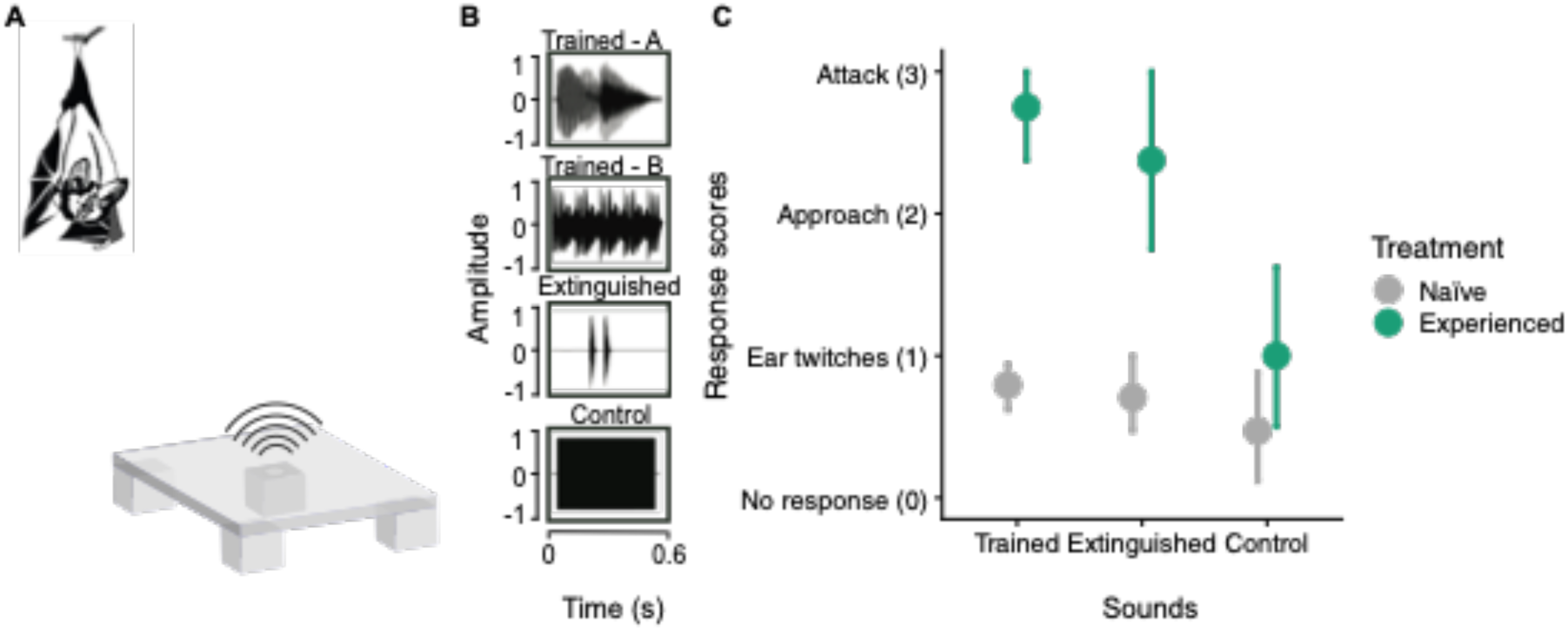
Experienced bats but not naïve bats attacked trained sounds. (A) Schematic of experimental setup (not to scale). (B) Oscillograms of experimental sounds. (C) Mean responses of naïve (gray) and experienced bats (green) with bootstrapped 95% confidence intervals.

## Discussion

Our results demonstrate remarkably long memories in wild frog-eating bats, with individuals remembering a learned foraging association for up to 4.2 years without reinforcement in the wild. This duration is comparable to that reported for corvids and primates (Table S1).The observation that six experienced bats also approached the previously extinguished sound (Fig. S1) suggests that either they remembered the difference between the sounds but resampled the extinguished sound, or they generalized the trained association to a sufficiently similar sound^7^. Previous work shows that frog-eating bats approach the calls of allopatric species that share acoustic characteristics with local palatable prey and avoid allopatric species’ calls that sound like local toxic prey^8^. Generalization over time may be adaptive given that older memories are less likely to reflect the current environment. When environments change, and especially when sampling costs are low, individuals may benefit from resampling^9^, and some sampling is necessary for trial-and-error learning. For example, three of the naïve bats approached novel sounds including the control sound, showing that bats occasionally investigate novel sounds.

Our study highlights that memory experiments with marked individuals at long-term field sites can help researchers link wild memory duration to species-specific ecological traits. Some of the preferred prey species of *T. cirrhosus* are either rare or explosive breeders that are heard infrequently during much of the year^10^. The ability for this bat to remember previously profitable prey cues over long time intervals would therefore allow them to avoid costly trial-and-error learning when exploiting these seasonal or rare resources. Comparative studies of cognitive ability across diverse taxa could be facilitated by leveraging the existence of marked wild individuals from long-term field studies.

### Ethical approval

All applicable international, national, and institutional guidelines for the care and use of animals were followed. All procedures performed in studies involving animals were in accordance with the ethical standards of the institution or practice at which the studies were conducted. Experimental procedures were licensed and approved by the Smithsonian Tropical Research Institute (IACUC permits: 20100816-1012-16 and 2014- 0101-2017) and the Autoridad Nacional del Ambiente de Panamá (SE/A-91-09, SE/A- 95-10, SE/A-6-11, SE/A-46-11, SE/A-94-11, SE/A-58-12, and SE/A-19-13).

## Acknowledgements

We would like to thank Dinielys Aparicio, Thomas Faughnan, Santiago Meneses and Michelle Simpson for assistance with the bats, and Katrine Hulgard and Inga Geipel for helpful comments on the manuscript. We are also grateful for an NSF GRFP and a short-term fellowship from the Smithsonian Tropical Research Institute to M. M. D.

## Author contributions

M. M. D., P. L. J, and R. A. P. designed the study. M. M. D. collected the data. M. M. D. and G. G. C. analyzed the results. M. M. D, P. L. J., M. M. R., and G. G. C. wrote and edited the manuscript.

## Supplemental material

### Supplemental data

Additional figures, methods, and discussion. (Below)

### Data and code

https://figshare.com/articles/dataset/LTM_frog_eating_bats_code_Rmd/19360397

### MP3 recording of the acoustic stimuli

https://figshare.com/articles/media/LTM_frog_eating_bats_Supplemental_data_acoustic_stimuli_mp3/19364693

### Video recordings showing responses of naïve and experienced bats

https://figshare.com/articles/media/LTM_frog_eating_bats_Supplemental_data_video_bat_responses_mov/19364717

## Supplemental data

**Table S1:**
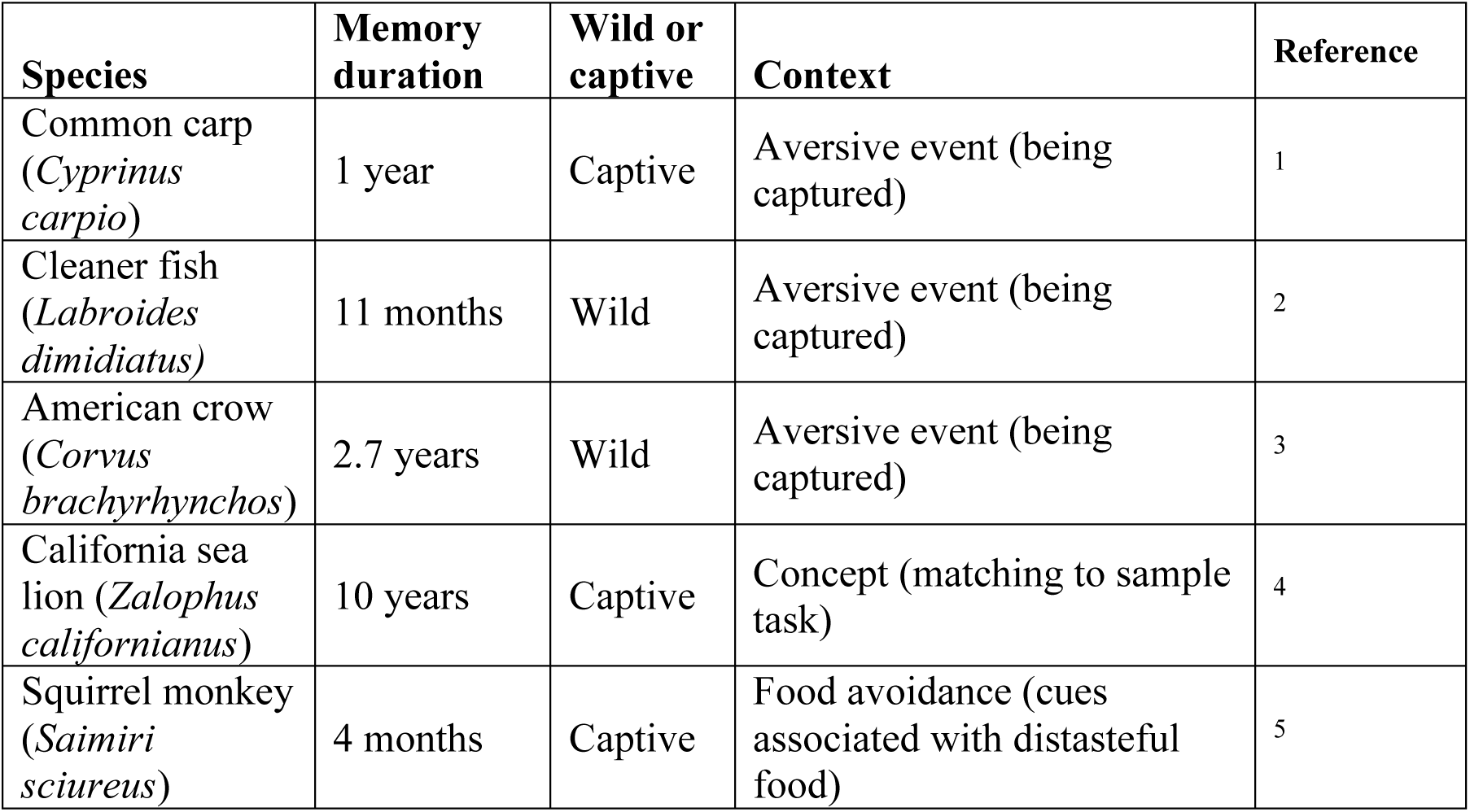
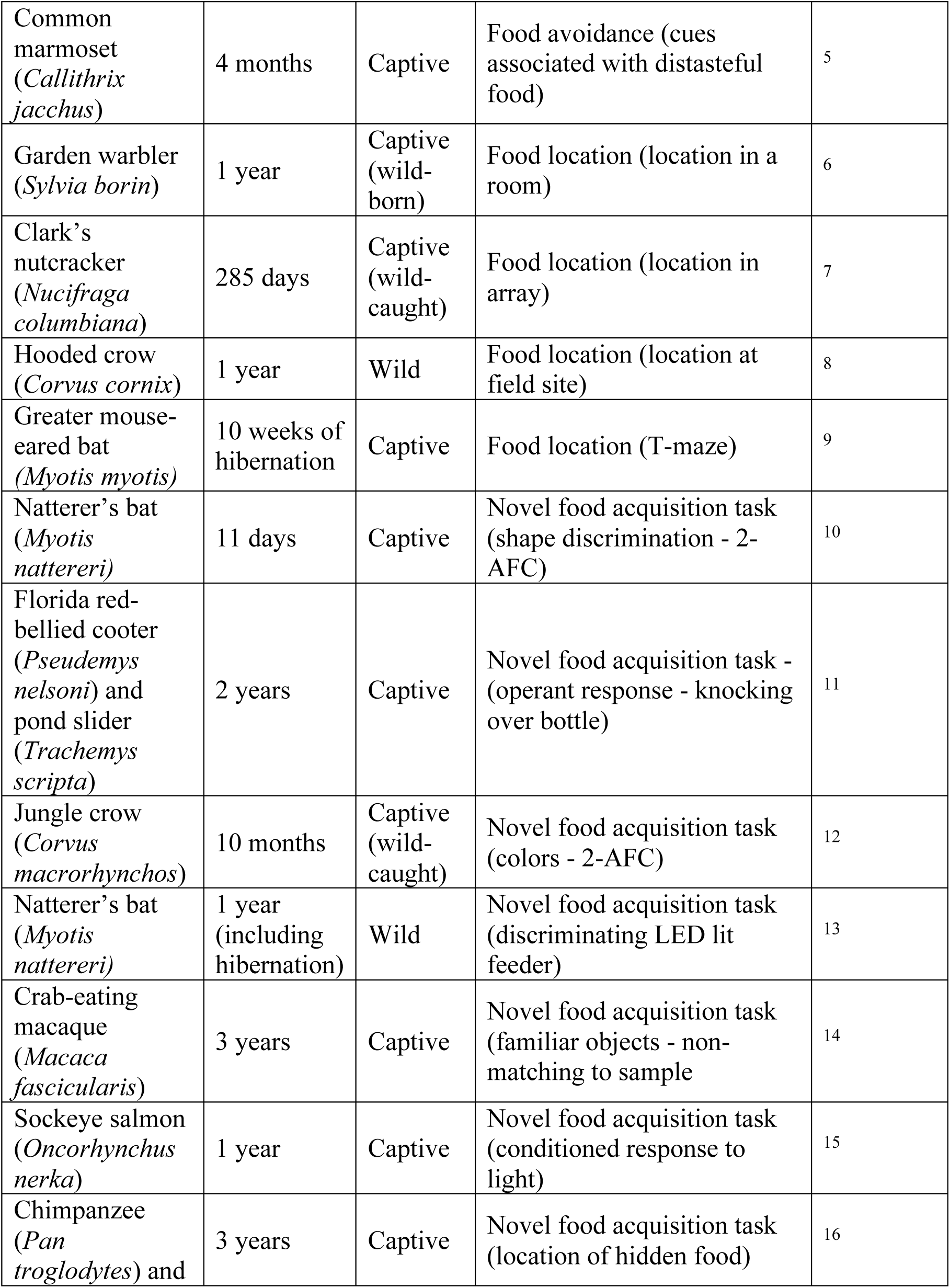
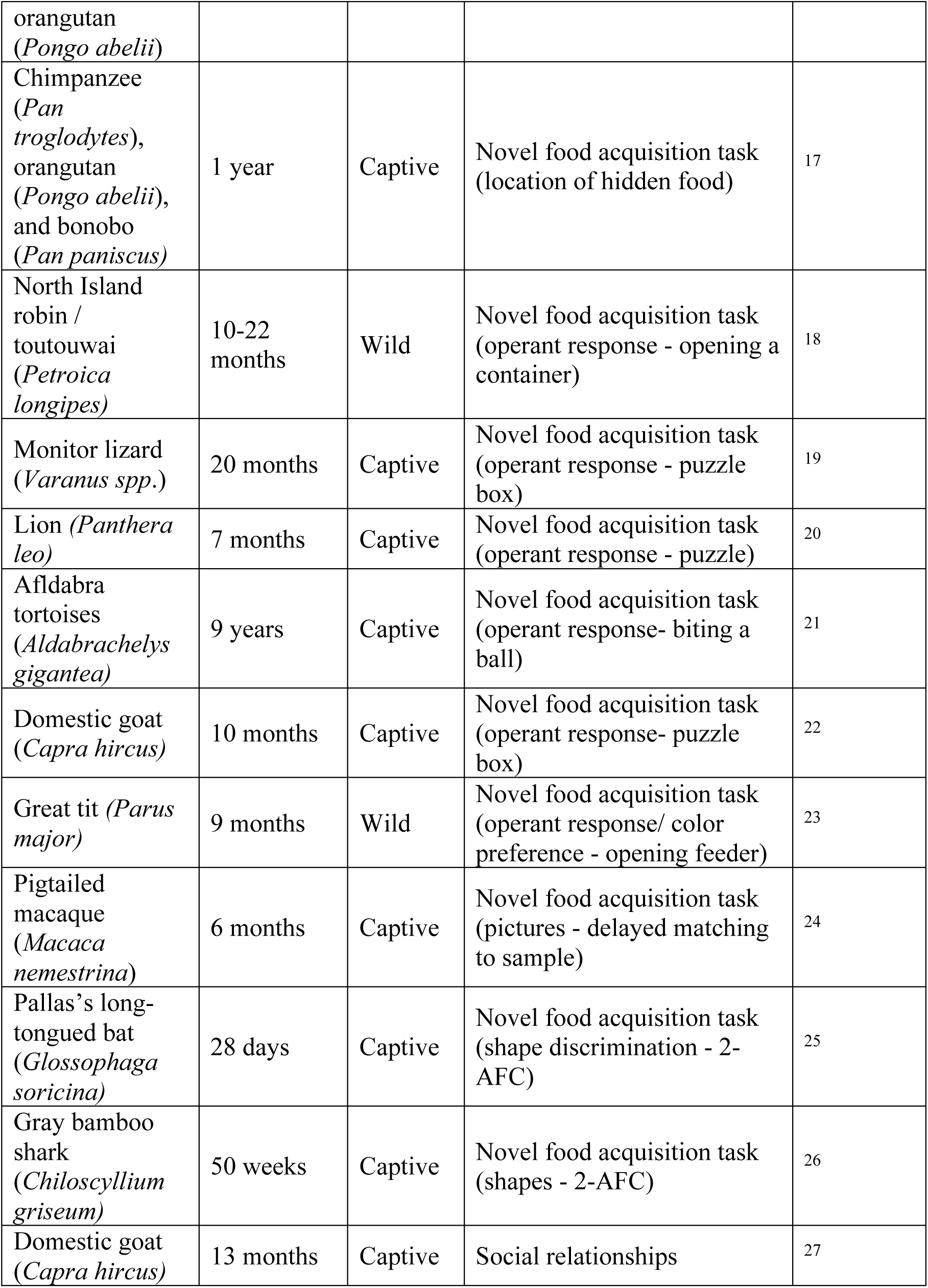
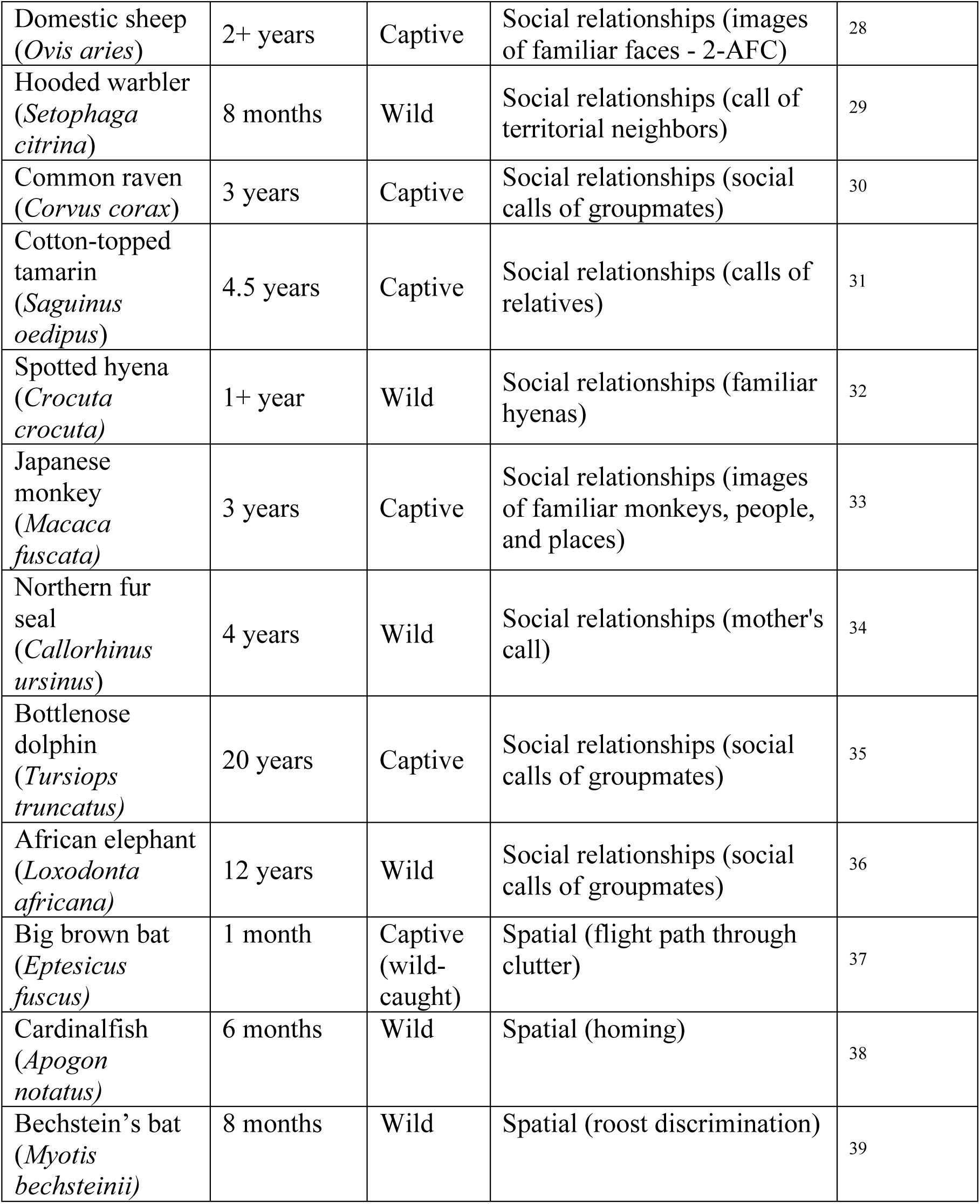
Studies of animal memory durations documenting memories of months or years. Memory duration = the longest duration demonstrated in the study. Wild or captive = where animals spent the time of memory retention; some “captive” animals were originally wild caught. Context = the ecological context and memory task being tested. 2-AFC = Two-alternative forced choice.

**Figure S1.**
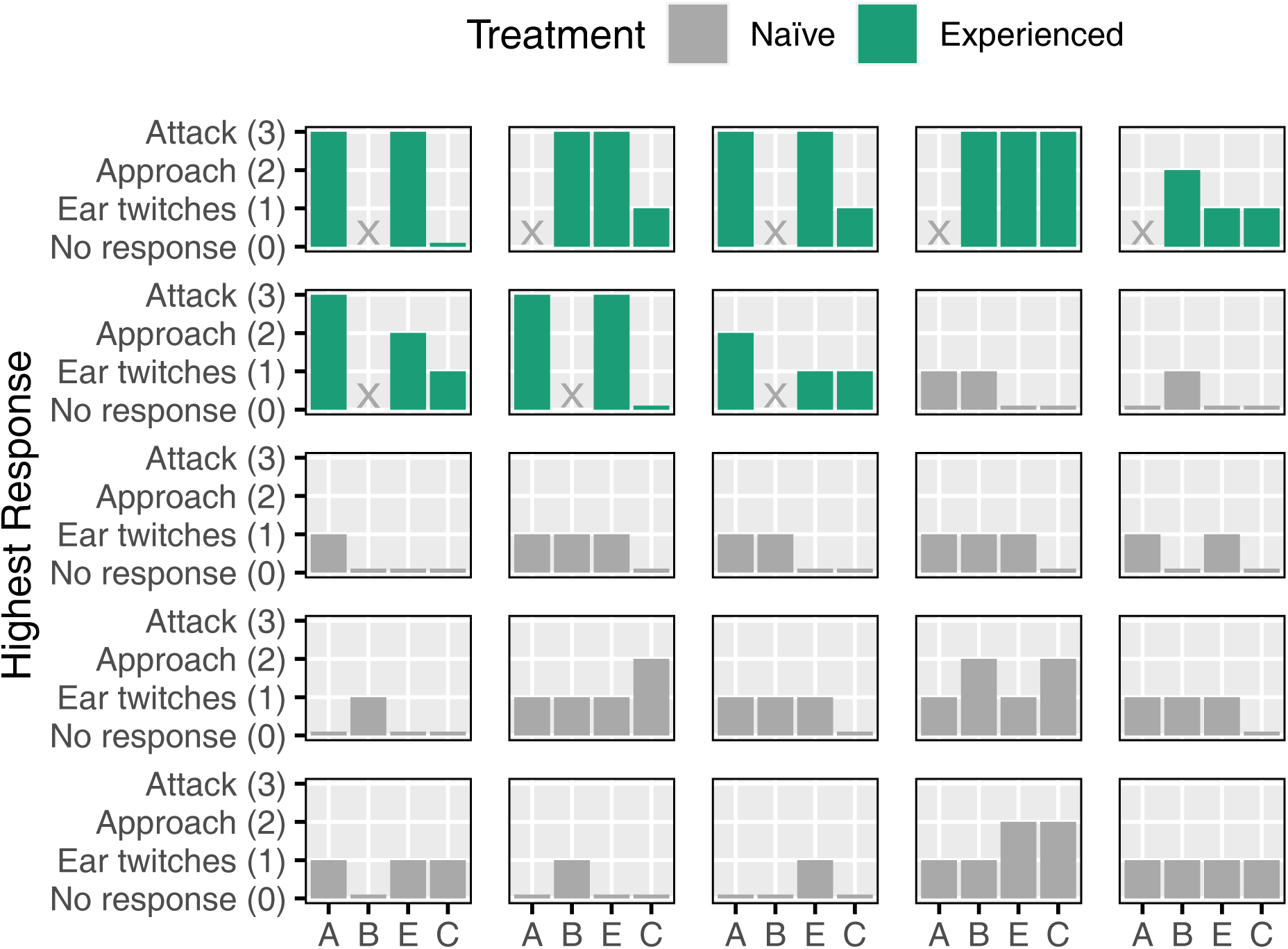
Experienced bats responded more than naïve bats. Highest responses of each bat to each sound. A = trained-A, B = trained-B, E = extinguished sound, C = control sound. Experienced bats are in green. Naïve bats are in gray. X indicates that stimulus was not tested. Number of days in the wild between training and retesting for the experienced bats (from left to right) was: 356, 444, 479, 501, 719, 818, 1524, 1531.

## Supplementary methods

We captured bats in mist nets at night and at their roosts during the day between February 2010 and July 2018 in Soberanía National Park, Panamá. Animals were maintained in a 5 × 5 × 2.5 m open-air flight cage at ambient temperature and humidity. All bats were uniquely marked with a subcutaneous passive integrated transponder (PIT tag, Trovan, Ltd.), and were released into the wild at their respective capture locations once they had been tested. Forty-nine bats were originally caught and trained to fly to a novel sound for food (cell phone ringtones) for a previous study (for complete methods, see^40^). Bats were initially lured to a speaker playing frog calls, where they learned to pick up a reward (pieces of baitfish) placed on a screen above the speaker. Once bats regularly attacked the speaker playing frog calls, they were trained to fly to a ringtone (either “trained-A” or “trained-B”) by gradually reducing the amplitude of the frog calls and playing the novel sounds progressively louder in successive steps. After learning to fly to their trained ringtone, bats spontaneously generalized the experimental association and would also fly to other ringtones. They were then given discrimination training, wherein rewarded trials playing their trained ringtone were interspersed with unrewarded trials with one of three other ringtones. Discrimination training quickly extinguished bats’ responses to the three unrewarded ringtones used in this study (means of 2.7 ± 0.9, 2.3 ± 0.8, and 3.3 ± 1.4 trials with each unrewarded sound to stop flying^40^).

We recaptured 8 of the 49 trained bats (hereafter ‘experienced’ bats) between January 2013 and July 2016, 356 to 1531 days after their initial release. Bats were tested under the same experimental conditions as the original training phase ^40^: on the first night after recapture we fed the bat baitfish by hand to acclimate it to captivity, and at the end of the night we released it into the flight cage. We began the experiment at the start of the second night. The experiment consisted of three trials, presented in random order. In each we played the bat one of three sounds: their trained ringtone, an extinguished ringtone, and a control sound (Fig. 1). The extinguished stimulus had similar acoustic properties to the trained ringtone in that it was broadband, frequency-modulated, and had the same call rate. The control sound was narrowband, not frequency- or amplitude-modulated, and was constant rather than intermittent (Fig. 1). All stimuli were broadcast at 80 dB SPL (re. 20 μPa) at 1m from the speaker. Ringtones were broadcast repeatedly at 2 s intervals, and the control sound was broadcast continuously, for a total of 30 s or until the bat flew to the speaker and retrieved a reward, whichever came first. Each trial began two minutes after the preceding trial or immediately after the bat finished eating. The 17 naïve bats had no prior experience with the experimental sounds. During testing, naïve bats were presented with all four sounds, including both trained-A and trained-B, to test for baseline responses to these stimuli.

Since we played naïve bats both trained-A and trained-B, while experienced bats were played only one or the other, we averaged each naïve bat’s responses to these two sounds to compare them to the experienced bat’s responses to the trained ringtone. Our conclusions do not change if we instead used the maximum of the two responses rather than the mean.

To test whether experienced bats had higher responses to the different sounds than the naïve bats, we used a non-parametric permutation test. To get a two-sided p-value, we compared the observed difference between the experienced and naïve groups in their average response to each sound to a distribution of 5000 values expected under the null hypothesis. We generated these values by randomly swapping the experienced vs naïve group label between the subjects and recalculating the mean difference in response between the groups 5000 times. To test whether the experienced bats responded differently to the different sounds, we ran a similar permutation test, where we calculated the average differences in the response to the trained, extinguished, and control sounds, and then compared those values against a null distribution created by randomly swapping the response scores between sounds within each bat. To generate 95% confidence intervals on response scores, we used nonparametric bootstrapping.

### Supplementary results and discussion

The experienced bats had stronger responses than the naïve bats to the trained ringtone (P < 0.0002) and extinguished ringtone (P < 0.0002), but not to the control sound (P = 0.2252), Fig. 1). Responses by individual bats can be seen in Fig. S1. Experienced bats responded weakly to the control sound (95% CI: 0.5 – 1.6), compared to the trained (95% CI: 2.4 – 3.0, P = 0.0004) and the extinguished ringtones (95% CI: 1.8 – 2.9, P = 0.0172). The naïve bats had similar responses to the four sounds (95% CIs: trained: 0.6 – 1.0, extinguished: 0.4 – 1.0, control: 0.1 – 0.9).

The difference between experienced and naïve bats is unlikely to be explained by age effects (older bats being more neophilic), or differences in comfort in captivity, for several reasons. Older bats in other species are less neophilic^41^, rather than more. One of the naïve bats in this study was previously captured 222 days before this experiment and had low responses (maximum response = 1). Likewise, in preliminary trials where naïve bats were tested without the control sound, four of the bats had been captured and tested in other experiments 101-1017 days previously, and these also showed low responses to the stimuli.

